# Metabolomic Profiling to Identify Early Urinary Biomarkers and Metabolic Pathway Alterations in Autosomal Dominant Polycystic Kidney Disease

**DOI:** 10.1101/2022.12.08.519365

**Authors:** Eden A. Houske, Matthew G. Glimm, Annika R. Bergstrom, Sally K. Slipher, Hope D. Welhaven, Mark C. Greenwood, Greta M. Linse, Ronald K. June, Alan S.L. Yu, Darren P. Wallace, Alyssa K. Hahn

**Affiliations:** Department of Biological and Environmental Science, Carroll College, Helena, MT, USA, 59625; Department of Chemical and Biological Engineering, Villanova University, Villanova, PA, USA, 19085; Department of Mathematical Sciences, Montana State University, Bozeman, MT, USA, 59717; Department of Chemistry and Biochemistry, Montana State University, Bozeman, MT, USA, 59717; Molecular Biosciences Program, Montana State University, Bozeman, MT, USA, 59717; Department of Mechanical and Industrial Engineering, Montana State University, Bozeman, MT, USA, 59717; Jared Grantham Kidney Institute, Department of Internal Medicine, University of Kansas Medical Center, Kansas City, KS

**Keywords:** Polycystic kidney disease, biomarkers, metabolomics, liquid chromatography-mass spectrometry, urine

## Abstract

Autosomal dominant polycystic kidney disease (ADPKD) is characterized by the formation of numerous fluid-filled cysts that lead to progressive loss of functional nephrons. Currently, there is an unmet need for diagnostic and prognostic indicators of early stages of the disease. Metabolites were extracted from the urine of early-stage ADPKD patients (n=48) and age- and sex-matched normal controls (n=47) and analyzed by liquid chromatography-mass spectrometry. Orthogonal partial least squares-discriminant analysis was employed to generate a global metabolomic profile of early ADPKD for the identification of metabolic pathway alterations and discriminatory metabolites as candidates of diagnostic and prognostic biomarkers. The global metabolomic profile exhibited alterations in steroid hormone biosynthesis and metabolism, fatty acid metabolism, pyruvate metabolism, amino acid metabolism, and the urea cycle. A panel of 46 metabolite features were identified as candidate diagnostic biomarkers. Notable putative identities of candidate diagnostic biomarkers for early detection include creatinine, cAMP, dCMP, various androgens (testosterone, 5alpha-androstane-3,17,dione, trans-dehydroandrosterone), betaine aldehyde, phosphoric acid, choline, 18-hydroxycorticosterone, and cortisol. Metabolic pathways associated with variable rates of disease progression included steroid hormone biosynthesis and metabolism, vitamin D3 metabolism, fatty acid metabolism, the pentose phosphate pathway, tricarboxylic acid cycle, amino acid metabolism, sialic acid metabolism, and chondroitin sulfate and heparin sulfate degradation. A panel of 41 metabolite features were identified as candidate prognostic biomarkers. Notable putative identities of candidate prognostic biomarkers include ethanolamine, C20:4 anandamide phosphate, progesterone, various androgens (5alpha-dihydrotestosterone, androsterone, etiocholanolone, epiandrosterone), betaine aldehyde, inflammatory lipids (eicosapentaenoic acid, linoleic acid, stearolic acid), and choline. Our exploratory data support metabolic reprogramming in early ADPKD and demonstrate the ability of liquid chromatography-mass spectrometry-based global metabolomic profiling to detect metabolic pathway alterations as new therapeutic targets and biomarkers for early diagnosis and tracking disease progression of ADPKD.

## Introduction

Autosomal dominant polycystic kidney disease (ADPKD) is the most common heritable kidney disease, affecting 1:400-1:1000 individuals worldwide(1). ADPKD is caused by mutations in *PKD1* or *PKD2*, which encode polycystin 1 (PC1) and polycystin 2 (PC2), respectively, and is characterized by the development and progressive enlargement of fluid-filled cysts in the kidneys(1). There is phenotypic variability in the clinical presentation of symptoms, severity of disease, rate of progression, and age of onset of end stage renal disease (ESRD)(2). Truncating mutations in *PKD1* are associated with a more severe phenotype (earlier symptom presentation, more and larger cysts, and faster rate of progression to renal failure) than non-truncating *PKD1* mutations or *PKD2* mutations(2). In all cases, renal cysts damage the surrounding tissue and ultimately lead to the decline in function(1).

ADPKD is often diagnosed by renal ultrasonography confirmation of cystic kidneys and a positive family history when patients present with symptoms such as abdominal and back pain, hypertension, urinary tract infections, hematuria, and kidney stones(1, 3, 4). Renal function and disease progression are then monitored using a variety of renal indicators including estimated glomerular filtration rate (eGFR) and quantification of plasma creatinine and blood urea nitrogen (BUN) levels and imaging techniques such as renal ultrasonography, computed tomography (CT), or magnetic resonance imaging (MRI)(3). However, the development of renal cysts may precede presentation of clinical symptoms and subsequent decline in renal function by many decades. Therefore, indicators of renal function are inadequate for detection or monitoring disease progression of early-stage ADPKD, signaling a critical need for improved methods for early diagnosis.

Current diagnostic methods do not determine the extent of damage to the kidneys due to early cyst formation or establish the rate of progression. Knowing which patients have aggressive disease would allow for selection of patients for clinical trials and treatment with Tolvaptan, the only FDA-approved treatment(5). MRI is the best modality for determining disease severity and rate of disease progression by monitoring height-adjusted total kidney volume (hTKV) relative to age – a prognostic model known as the Mayo Imaging Classification (MIC)(6). However, MRI is expensive and requires multiple scans over several months to years to determine changes. Thus, there is a need for new inexpensive and minimally invasive methods for diagnosing and monitoring disease progression of early-stage ADPKD to determine eligibility to receive new therapeutic interventions to slow or halt the progression toward ESRD.

An emerging area in ADPKD research is understanding the role of altered cellular metabolism in cystogenesis. Recent work suggests that ADPKD is associated with profound metabolic reprogramming as a key driver of cystogenesis(7, 8). Considering that early in ADPKD, dilated cystic structures are still in contact with the collecting system and are likely excreted into the urine, profiling the urinary metabolome may reveal novel biomarkers and the discovery of perturbed metabolic pathways associate with early cystogenesis. Only a handful of studies have analyzed urine from ADPKD patients in search of metabolic biomarkers of ADPKD(9-11). These studies reported candidate diagnostic biomarkers including lactate, pyruvate, succinate, cyclic AMP (cAMP), myo-inositol, creatinine, 3-hydroxyisovalerate, asymmetric dimethylarginine, the ratio of alanine over citrate, formate, tartaric acid, citrate, threonine, methanol, sucrose, alanine, 6-hydroxynicotinic acid, D-saccharate, and tyrosine(9-11). Two recent studies that analyzed the urinary metabolome in patients with ADPKD also correlated metabolic markers with current indicators of renal function decline (eGFR, hTKV) to identify prognostic biomarkers of ADPKD(9, 11). Of the candidate biomarkers listed above, lactate, pyruvate, succinate, and cAMP correlated with disease severity (lower eGFR, higher hTKV) and the alanine/citrate ratio was most strongly associated with faster rate of disease progression (annual change in eGFR)(9, 11).

The current study is focused on changes in urinary metabolic biomarkers in early-stage ADPKD, before irreversible damage to the kidneys and decline in function. The early-stage ADPKD patient population in this study is younger and exhibits greater preservation of renal function in comparison to the previous studies reported above. To our knowledge, this is the first study to analyze the urinary metabolome for prognostic metabolic markers related to MIC subclasses, which is the current prognostic model used in clinical settings to predict the rate of disease progression. Thus, the overall goal is to expand our understanding of altered cellular metabolism in ADPKD toward the development of new diagnostics for earlier detection and new prognostic markers. We accomplished this by analyzing urinary metabolite extracts from a cohort of early-stage ADPKD patients with preserved renal function by liquid chromatography-mass spectrometry (LC-MS) to generate a global metabolomic profile of early-stage ADPKD in search of diagnostic and prognostic indicates of early disease.

## Methods

### Sample population

Urine samples were obtained from the PKD Biomarkers and Biomaterials Core in the Kansas PKD Research and Translation Core Center (U54), which is part of the National PKD Research Resource Consortium. The Core repository contains urine samples obtained from the longitudinal Early PKD Observational Cohort (EPOC) study under Institutional Review Board (IRB) approval. The EPOC study collects urine from study participants with early-stage ADPKD (ages 4 to 35 years at enrollment) with clinical data including age, sex, plasma creatinine, BUN, eGFR, htTKV, race, body mass index (BMI), and subject relationship if any, along with age-matched healthy controls (Table 1). ADPKD patients were classified according to the MIC into subclasses 1A-1E (ranked in order from slowest progression to most rapid) based on hTKV relative to age as a prognostic model of rate of disease progression (Table 2). Adult participants were asked to fast for 8 hours and children for 4 hours prior to the study visit and second morning urines were collected. Urine samples were obtained from 48 study participants with early-stage ADPKD at baseline and 47 healthy normal controls.

**Table 1.** Clinical characteristics of ADPKD and control cohorts. Study participants are classified as control or ADPKD cohorts. Cohorts are reported with percentage of males and females and mean (with standard deviation; SD) age, body mass index (BMI), estimated glomerular filtration rate (eGFR), blood urea nitrogen (BUN) levels, creatinine levels, and height-adjusted total kidney volume (hTKV) if obtained with study participant samples. hTKV measurements were not obtained for control study participants.

**Clinical Characteristics (ADPKD and Control Groups)**
|  | Control (n = 47) | ADPKD (n = 48) | p-value<sup>1</sup> |
| --- | --- | --- | --- |
| **Sex, n (%)** |  |  | 0.12 |
| Female | 25 (53%) | 33 (69%) |  |
| Male | 22 (47%) | 15 (31%) |  |
| **Age (years), Mean (SD)** | 22 (6) | 23 (6) | 0.68 |
| **BMI (kg/m<sup>2</sup>), Mean (SD)<sup>2</sup>** | 25.1 (6.6) | 24.7 (4.5) | 0.65 |
| **BUN (mg/dL), Mean (SD)** | 13 (4) | 13 (4) | 0.35 |
| **Creatinine (mg/dL), Mean (SD)** | 0.81 (0.21) | 0.80 (0.18) | 0.88 |
| **eGFR (mL/min/1.73m<sup>2</sup>), Mean (SD)** | 116 (21) | 113 (19) | 0.59 |
| **hTKV (mL/m), Mean (SD)** |  | 491 (364) |  |
| <sup>1</sup> Pearson's Chi-squared test; Wilcoxon rank sum test |  |  |  |
| <sup>2</sup> 1 control group subject unknown (excluded from BMI summary) |  |  |  |

**Table 2.** Clinical characteristics of MIC subclasses. ADPKD study participants are separated into MIC subclasses 1A-1E and are reported with percentage of males and females and mean (with standard deviation; SD) age, body mass index (BMI), estimated glomerular filtration rate (eGFR), blood urea nitrogen (BUN) levels, creatinine levels, height-adjusted total kidney volume (hTKV), and mutation.

| Clinical Characteristics (Mayo Imaging Classification Subclasses) |  |  |  |  |  |  |
| --- | --- | --- | --- | --- | --- | --- |
|  | 1A (n = 6) | 1B (n = 8) | 1C (n = 15) | 1D (n = 6) | 1E (n = 13) | p-value<sup>1</sup> |
| **Sex, n (%)** |  |  |  |  |  | 0.21 |
| Female | 6 (100%) | 6 (75%) | 11 (73%) | 4 (67%) | 6 (46%) |  |
| Male | 0 (0%) | 2 (25%) | 4 (27%) | 2 (33%) | 7 (54%) |  |
| **Age (years), Mean (SD)** | 20 (5) | 26 (5) | 25 (5) | 25 (5) | 20 (6) | 0.029 |
| **BMI (kg/m<sup>2</sup>), Mean (SD)** | 22.0 (4.4) | 23.8 (4.3) | 24.4 (4.0) | 25.9 (3.6) | 26.1 (5.4) | 0.43 |
| **BUN (mg/dL), Mean (SD)** | 12 (2) | 14 (5) | 12 (3) | 13 (2) | 16 (5) | 0.19 |
| **Creatinine (mg/dL), Mean (SD)** | 0.71 (0.08) | 0.83 (0.16) | 0.81 (0.18) | 0.79 (0.23) | 0.83 (0.22) | 0.79 |
| **eGFR (mL/min/1.73m<sup>2</sup>), Mean (SD)** | 116 (11) | 107 (14) | 111 (23) | 114 (20) | 116 (22) | 0.73 |
| **hTKV (mL/m), Mean (SD)** | 210 (45) | 270 (32) | 380 (98) | 570 (152) | 847 (519) | <0.001 |
| **Mutation, n (%)<sup>2</sup>** |  |  |  |  |  | 0.005 |
| PKD1 | 2 (40%) | 7 (100%) | 13 (93%) | 6 (100%) | 13 (100%) |  |
| PKD2/NMD | 3 (60%) | 0 (0%) | 1 (7.1%) | 0 (0%) | 0 (0%) |  |

<sup>1</sup> Fisher's exact test; Kruskal-Wallis rank sum test
<sup>2</sup> 2 subjects not tested, 1 subject with PKD1 and PKD2 (excluded from mutation summary)

### Metabolite extraction

Metabolites were extracted from urine using an established metabolite extraction protocol for human biofluids, with slight modifications/adaptations for urine(12-16). 100 uL aliquots of urine were thawed on ice, and four volumes of ice-cold methanol (-20°C) were added for a single, combined protein precipitation and metabolite extraction step. Samples were vortexed for 2 min and subsequently placed at -20°C for 30 min. Samples were subjected to centrifugation at 16,100 x *g* for 10 min at 4°C. Supernatants were collected and dried in a vacuum concentrator for 2 hr. Dried metabolite extracts were frozen at -80°C until mass spectrometry analysis.

### Liquid chromatography-mass spectrometry analysis

Dried metabolite extracts were resuspended in mass spectrometry grade 50:50 water: acetonitrile. For quality control purposes, an injection blank containing mass-spectrometry grade water and a pooled sample with 1-2 uL from each study sample were created. Both quality control samples were injected and analyzed multiple times throughout the batch analysis to assess LC-MS system performance. Samples were analyzed in positive mode using an Agilent 1290 UPLC system coupled to an Agilent 6538 Quadrupole-Time of Flight (Q-TOF) mass spectrometer (Agilent, Santa Clara, CA). Metabolites were separated on a Cogent Diamond Hydride HILIC column (150 × 2.1 mm, MicroSolv, Eatontown, NJ) using an optimized normal phase 25-minute gradient elution method(12-14). Mass spectra were processed as previously described(12).

### Statistical analysis

Metabolite features with a relative abundance of zero were either considered non-detects or low detection (below noise threshold) and therefore were replaced with one-fifth the minimum peak intensity. Prior to statistical analyses, data were normalized using a modified probabilistic quotient normalization (PQN) to the QC pooled samples to account for varying dilution factors in complex biofluids that could result in dramatic variations in overall concentrations of metabolites in urine(17). Data were log transformed using the base-10 logarithm (log10) to help to correct for non-normal distributions commonly observed in these data and then standardized (mean centered divided by standard deviation). The dataset processed as described above was used for all further statistical analyses unless otherwise indicated.

Orthogonal partial least squares discriminant analysis (OPLS-DA) was used in search of differences in metabolomic profiles between cohorts and MIC subclasses(18, 19). Fit and reliability of models were assessed based on R^2^ (variation explained in metabolites, X, and classes, Y) and Q^2^ (predictive accuracy) values, with values closer to 1 suggesting better performance. Because OPLS-DA models are prone to overfitting, the models were further assessed with permutation tests of 1000 permutations. Importantly, permutation testing was considered a better indicator of model quality because OPLS-DA model metrics for assessing model fit were originally developed for quantitative, not qualitative categorical models, and permutation testing directly indicates how well the model classifies as compared to random chance (smaller p-values suggest performance that is better than chance). To assess for potentially confounding clinical covariates, statistical tests for group differences were performed, and OPLS-DA scores plots were coded by clinical characteristics and visually inspected.

Metabolite features with the greatest ability to discriminate between (1) control and ADPKD cohorts and (2) MIC subclasses were selected as important according to the “elbow” in the scree plot of Variable Importance in Projection (VIP) scores. The top discriminatory features were selected as candidate biomarkers of ADPKD for early detection and/or determining rate of disease progression. Metabolite features identified as candidate biomarkers of interest (*m/z* values) were matched to putative metabolite identities using the metabolite mass spectral database METLIN (mass tolerance: 30 ppm, exclude drugs, toxins, halogens, peptides, limited to metabolites with KEGG and CAS IDs)(20). Metabolite features with the same *m/z* values may match to multiple putative metabolite identities.

### Pathway analysis

To gain insight into metabolic shifts detected in urine early in disease progression, metabolite features with the greatest ability to discriminate between (1) control and ADPKD cohorts and (2) MIC subclasses with an OPLS-DA VIP score ≥ 1 were mapped to metabolic pathways using the Functional Analysis module in the metabolomics online program, MetaboAnalyst(21). A more lenient VIP score threshold was selected to enhance the performance of the pathway enrichment analysis algorithm which has limits for the minimum number of important metabolites to map. The Functional Analysis module uses the *mummichog* algorithm to predict a network of functional activity based on the projection of detected metabolite features onto local pathways. Pathway library Human MFN was used for compound identification and pathway enrichment (mass tolerance: 5ppm; positive mode, retention time consideration). Pathways are reported as significant by pathway overrepresentation analysis with an adjusted *a priori* significance level of 0.05 based on the gamma distribution to control for multiple tests.

## Results

### ADPKD Patient Characteristics

A total of 48 ADPKD patients, mean age of 22 (SD = 6) years and mean BMI of 24.7 (SD = 4.5) kg/m^2^, were included in this study. The ADPKD patient population had a mean eGFR of 113 (SD = 19) ml/min per 1.73 m^2^, plasma creatinine levels of 0.80 (SD = 0.18) mg/dL, and BUN levels of 13 (SD = 4) mg/dL, indicative of normal renal function. The ADPKD patient population had a hTKV of 491 (SD = 364) ml/m at the time of enrollment, consistent with early stages of disease. There were no differences in eGFR or levels of plasma creatinine or BUN between the ADPKD patient cohort and control study participants. Thus, our ADPKD cohort had preserved renal function and mild cystic kidney disease(22). Notably, there were no significant differences between the ADPKD and control study participant in other clinical characteristics (Table 1), reducing the possibility of potentially confounding clinical co-variates.

### Metabolic Differences Detected Between Control and ADPKD Cohorts

A total of 1554 metabolite features were detected across all 95 samples. Global metabolomic profiles of control and ADPKD cohorts were visualized by OPLS-DA in search of differences between cohorts. OPLS-DA showed separation of the majority of control and ADPKD study participants between their respective cohorts (Fig. 1). The OPLS-DA model had an 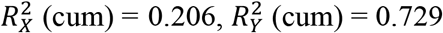, and Q^2^(cum) = -0.002. A permutation test with 1000 permutations resulted in p = 0.182 for 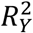 and p = 0.017 for Q^2^.

**Figure 1.**
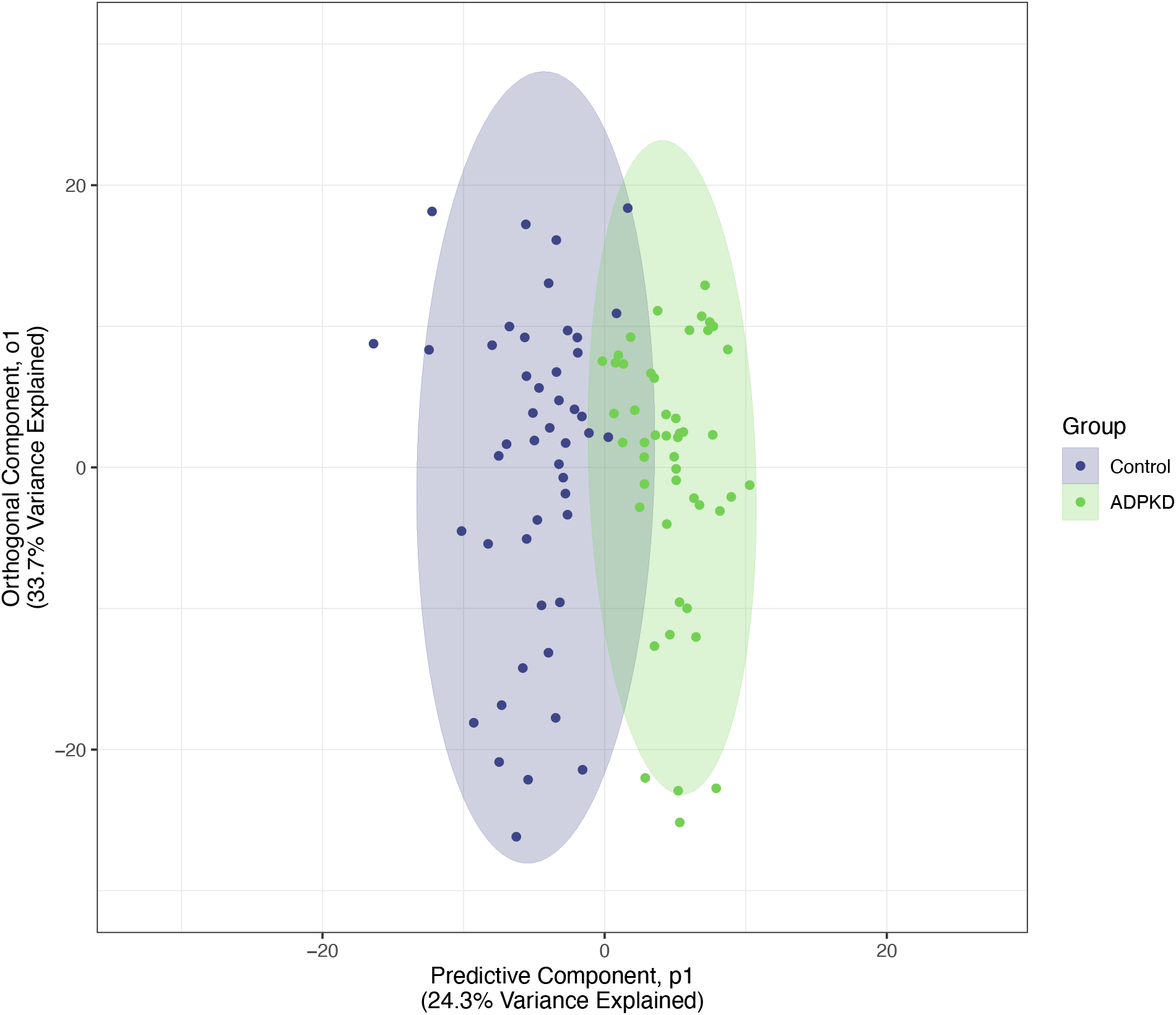
OPLS-DA scores plot shows separation between ADPKD and control study participants. OPLS-DA scores plot of ADPKD (green) and control (blue) samples. The predictive component p1 (x-axis) showcases inter-group variability. The orthogonal component o1 (y-axis) showcases intra-group variability. The OPLS-DA scores plot shows separation of the majority of study participants between their respective cohorts.

Metabolite features with the greatest ability to discriminate between control and ADPKD cohorts were identified based on OPLS-DA VIP scores. 573 metabolite features had a VIP score ≥ 1, indicating their ability to best discriminate between control and PKD cohorts. To narrow this list of important metabolite features further, 46 features with a VIP score ≥ 1.91, the elbow of the VIP scree plot, were selected as candidate biomarkers for early detection of ADPKD (Fig. 2A-B; Supplemental Fig. 2; Supplemental Table 1). Notable putative identities of candidate diagnostic biomarkers for early detection include creatinine, cAMP, dCMP, various androgens (testosterone, 5alpha-androstane-3,17,dione, trans-dehydroandrosterone), betaine aldehyde, phosphoric acid, choline, 18-hydroxycorticosterone, and cortisol (Supplemental Table 1).

**Figure 2.**
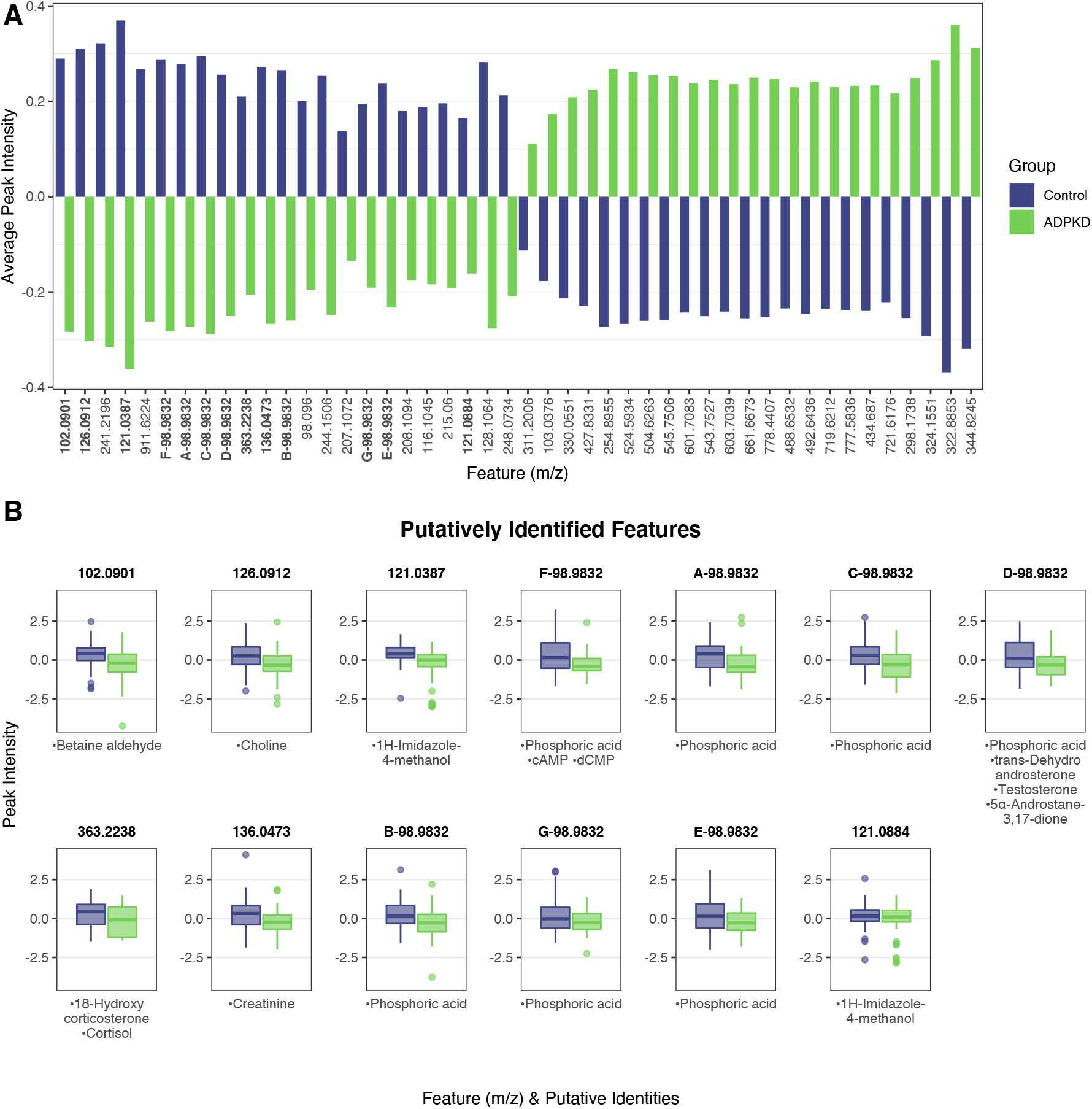
Important metabolite features between control and ADPKD cohorts selected as candidate diagnostic biomarkers. **(A)** Bar graph of mean log transformed peak intensities of important metabolite features selected from scree plot based on VIP score threshold ≥ 1.9. Features are ordered by hierarchical clustering on group averages. **(B)** Box plots of log transformed peak intensities of select candidate diagnostic biomarkers with matched KEGG IDs. Putative identities and corresponding *m/z* values are reported. For both (A) and (B), features named as A-, B-, etc. represent metabolite features with the same *m/z* value but distinct retention times. Blue represents the control cohort and green represents the ADPKD cohort. The full list of important features selected as candidate diagnostic biomarkers with matched putative identities is reported in Supplemental Table 3.

To generate a global metabolomic profile of metabolic shifts early in disease progression, metabolite features with the greatest ability to discriminate between control and ADPKD cohorts with an OPLS-DA VIP score ≥ 1 were mapped to metabolic pathways. These metabolite features that had the best ability to discriminate between control and ADPKD cohorts mapped to 14 significant pathways including androgen and estrogen biosynthesis and metabolism, steroid hormone biosynthesis and metabolism, the carnitine shuttle, fatty acid metabolism, pyruvate metabolism, amino acid metabolism (lysine and tyrosine), and the urea cycle (FDR-adjusted p-value < 0.05; Table 3; Supplemental Table 2).

**Table 3.**
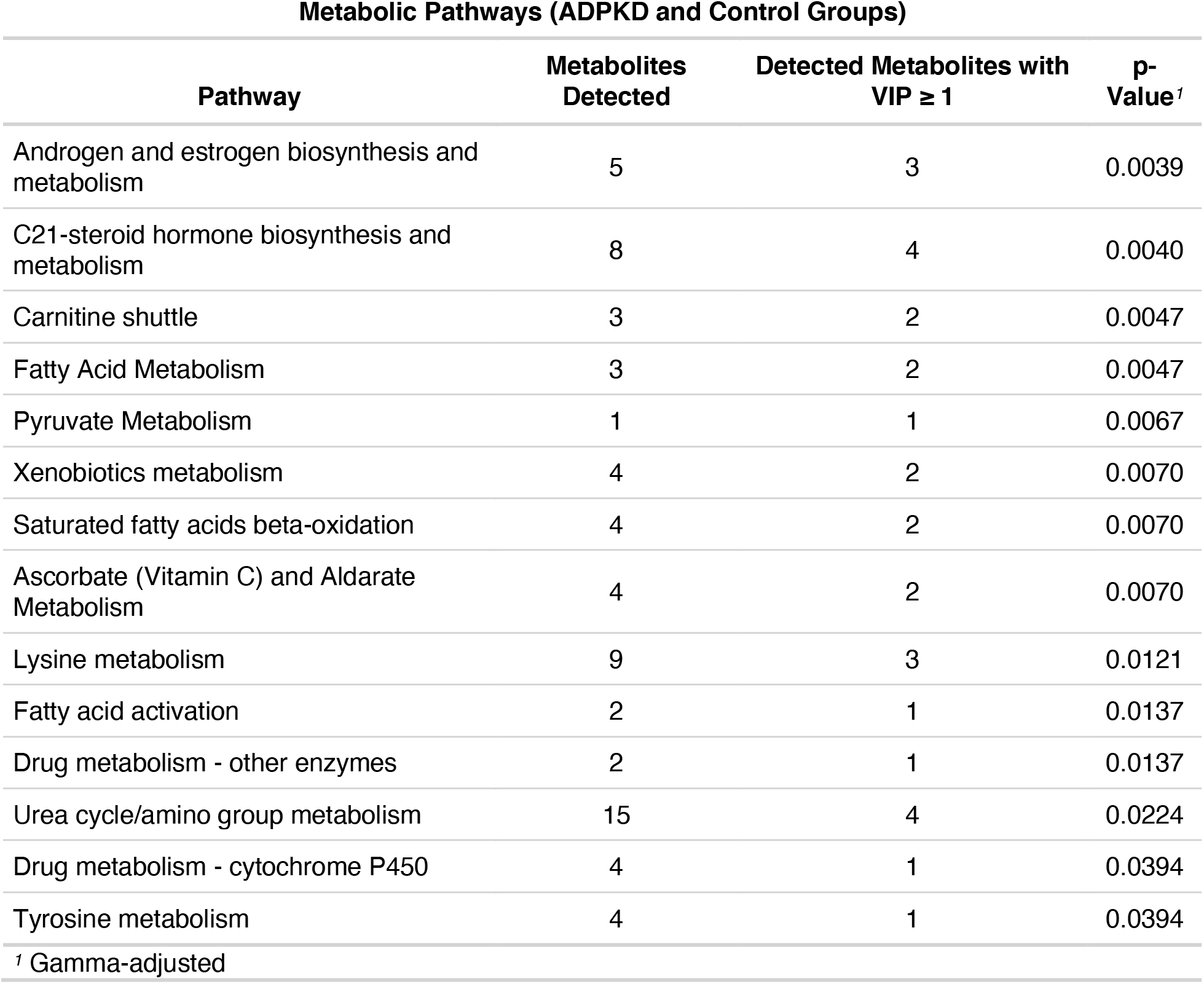
Metabolic pathways altered in early-stage ADPKD. Metabolite features with the greatest ability to discriminate between ADPKD and control study participants in OPLS-DA (VIP score ≥ 1) were mapped to metabolic pathways using the Functional Analysis module in the metabolomics online program, MetaboAnalyst. Pathways are reported with the total number of metabolite features detected in that pathway, total number of metabolites detected in the pathway with a VIP score ≥ 1, and Gamma-adjusted p-value for the pathway. The full list of altered metabolic pathways is reported in Supplemental Table 2.

Potentially confounding clinical covariates were further assessed through visualization of OPLS-DA scores plots coded by each covariate, with no noticeable differences observed (Supplemental Fig. 1).

### ADPKD Cohort Characteristics based on Mayo Imaging Classification

ADPKD participants were stratified using the MIC subclasses 1A-1E (ranked in order from slowest progression to most rapid), which uses hTKV relative to age as a prognostic model of rate of disease progression (Table 2). There were no statistically significant differences in sex, BMI, BUN levels, plasma creatinine levels, or eGFR among the subclasses. However, there were statistically significant differences in age, hTKV, and PKD gene mutation (*PKD1* vs. *PKD2/NMD*) between MIC subclasses.

### Metabolic Differences Detected in ADPKD Cohort Depending on Mayo Imaging Classification

OPLS-DA was used to visualize global metabolomic profiles of MIC subclasses in search of metabolic differences associated with different rates of disease progression (Fig. 3). OPLS-DA scores plot revealed some separation among the MIC subclasses 1A-1E, with the predictive component scores increasing with more rapid disease progression (Fig. 3). The OPLS-DA model had and 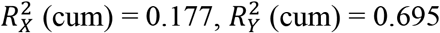, Q^2^(cum) = -0.263. A permutation test with 1000 permutations resulted in p = 0.645 for 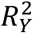 and p = 0.503 for Q^2^. The clearest separation was exhibited between subclasses with the slowest rate of disease progression (1A-B) and fastest rate of disease progression (1D-E), with subclass 1C exhibiting overlap between 1A-B and 1D-E. These results suggest that distinct metabolic phenotypes of ADPKD exist that correlate with rate of disease progression and thus may have unique prognostic metabolic biomarkers.

**Figure 3.**
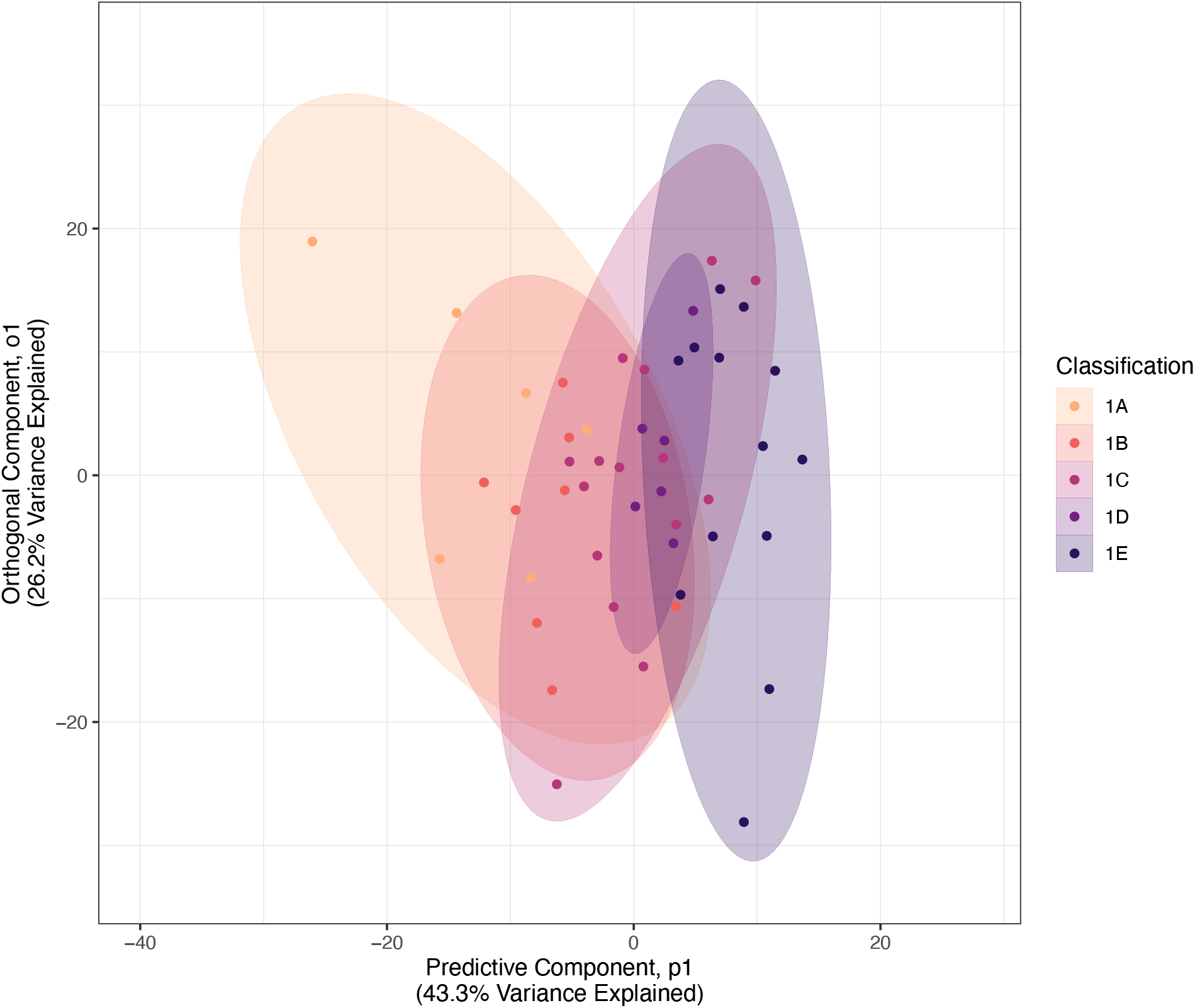
OPLS-DA scores plot of MIC subclasses shows separation between subclasses with slow and fast rates of disease progression. The OPLS-DA scores plot exhibits some separation among the MIC subclasses. The predictive component p1 (x-axis) showcases inter-group variability. The orthogonal component o1 (y-axis) showcases intra-group variability.

Important metabolite features with the greatest ability to discriminate between MIC subclasses were identified based on OPLS-DA VIP scores. 564 metabolite features had a VIP ≥ 1, indicating their ability to best discriminate between MIC subclasses. To narrow this list of important metabolite features further, 41 metabolite features were selected as candidate prognostic biomarkers according to a VIP score threshold ≥ 2, representative of the VIP score scree plot elbow (Fig. 4A-B, Supplemental Fig. 4, Supplemental Table 3). Notable putative identities of candidate prognostic biomarkers include ethanolamine, C20:4 anandamide phosphate, progesterone, various androgens (5alpha-dihydrotestosterone, androsterone, etiocholanolone, epiandrosterone), betaine aldehyde, inflammatory lipids (eicosapentaenoic acid, linoleic acid, stearolic acid), and choline (Supplemental Table 3).

**Figure 4.**
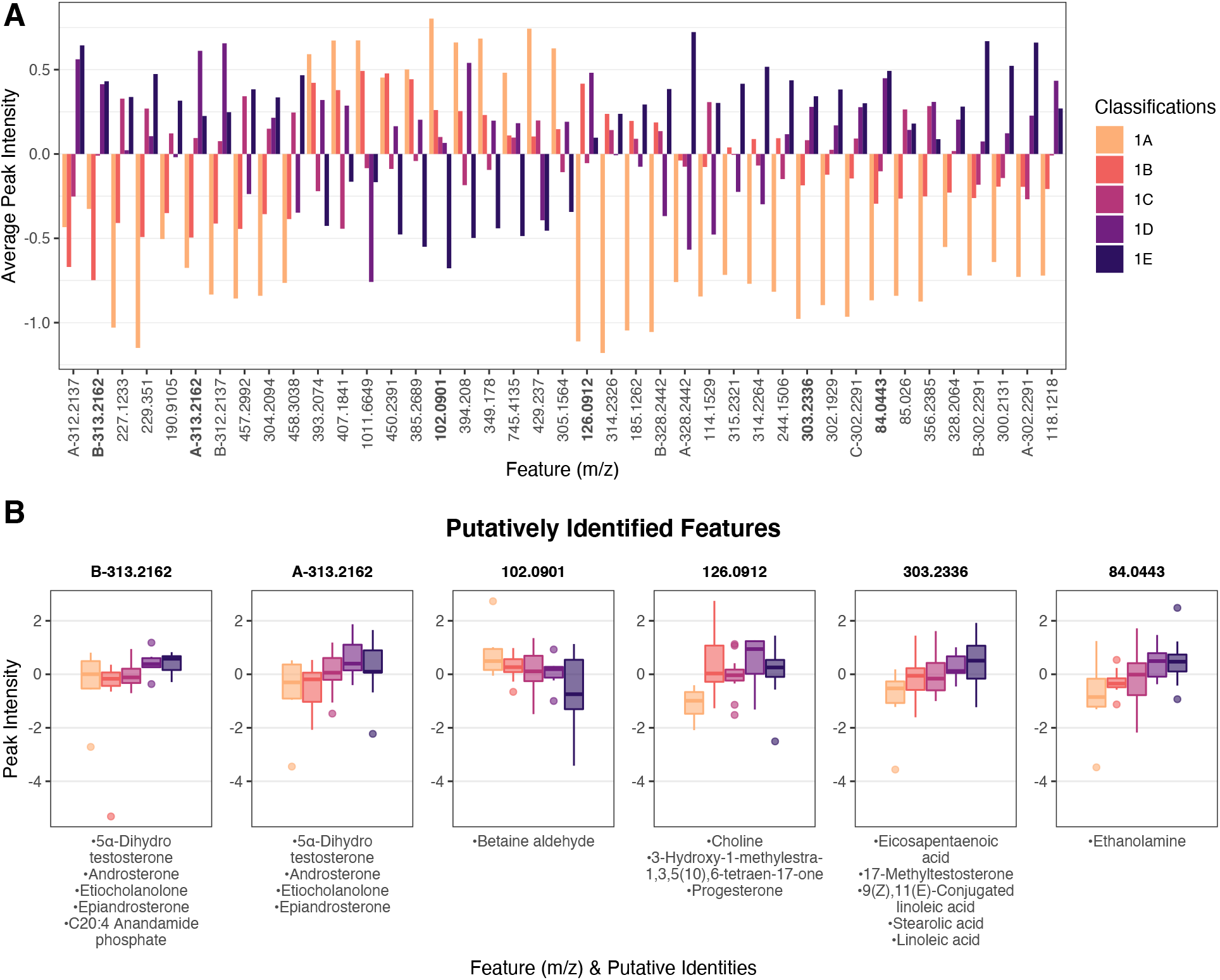
Important metabolite features between MIC subclasses selected as candidate prognostic biomarkers. **(A)** Bar graph of mean log transformed peak intensities of important metabolite features selected from scree plot based on VIP score threshold ≥ 2. Features are ordered by hierarchical clustering on group averages. **(B)** Box plots of log transformed peak intensities of select candidate diagnostic biomarkers with matched KEGG IDs. Putative identities and corresponding *m/z* values are reported. For both (A) and (B), features named as A-, B-, etc. represent metabolite features with the same *m/z* value but distinct retention times. MIC subclasses are colored by decreasing intensity (1A – light orange; 1E dark purple). The full list of important features selected as candidate prognostic biomarkers with matched putative identities is reported in Supplemental Table 3.

To gain insight into metabolic shifts associated with rate of disease progression, metabolite features with the greatest ability to discriminate between MIC subclasses based on an OPLS-DA VIP score ≥ 1 were mapped to metabolic pathways. Metabolite features with the greatest ability to discriminate between the MIC subclasses mapped to 38 significant pathways including steroid hormone biosynthesis and metabolism (in particular, androgen and estrogen), vitamin D3 metabolism, a variety of fatty acid-related pathways (de novo fatty acid biosynthesis, arachidonic acid metabolism, carnitine shuttle, linoleate metabolism, fatty acid oxidation, leukotriene metabolism, glycerophospholipid and glycosphingolipid metabolism, etc.), the pentose phosphate pathway (PPP), tricarboxylic acid cycle (TCA) cycle, amino acid metabolism (methionine, cysteine, arginine, proline, glutamate, histidine, tryptophan, and lysine), sialic acid metabolism, and chondroitin sulfate and heparin sulfate degradation (FDR-adjusted p-value < 0.05; Table 4; Supplemental Table 4).

**Table 4.** Metabolic pathway alterations that associate with variable rates of disease progression. Metabolite features with the greatest ability to discriminate between MIC subclasses in the OPLS-DA (VIP score ≥ 1) were mapped to metabolic pathways using the Functional Analysis module in the metabolomics online program, MetaboAnalyst. Pathways are reported with the total number of metabolite features detected in that pathway, total number of metabolites detected in the pathway with a VIP score ≥ 1, and Gamma-adjusted p-value for the pathway. The full list of altered metabolic pathways is reported in Supplemental Table 4.

| Metabolic Pathways (Mayo Imaging Classification Subclasses) |  |  |  |
| --- | --- | --- | --- |
| Pathway | Metabolites Detected | Detected Metabolites with VIP $\geq 1$ | p-Value<sup>†</sup> |
| C21-steroid hormone biosynthesis and metabolism | 8 | 8 | 0.0022 |
| Androgen and estrogen biosynthesis and metabolism | 5 | 4 | 0.0042 |
| Vitamin D3 (cholecalciferol) metabolism | 2 | 2 | 0.0054 |
| De novo fatty acid biosynthesis | 2 | 2 | 0.0054 |
| Arachidonic acid metabolism | 2 | 2 | 0.0054 |
| Drug metabolism - cytochrome P450 | 4 | 3 | 0.0068 |
| Ascorbate (Vitamin C) and Aldarate Metabolism | 4 | 3 | 0.0068 |
| Sialic acid metabolism | 1 | 1 | 0.0146 |
| Chondroitin sulfate degradation | 1 | 1 | 0.0146 |
| Linoleate metabolism | 1 | 1 | 0.0146 |
| Carnitine shuttle | 3 | 2 | 0.0146 |
| Parathio degradation | 1 | 1 | 0.0146 |
| Vitamin B9 (folate) metabolism | 1 | 1 | 0.0146 |
| Glyoxylate and Dicarboxylate Metabolism | 1 | 1 | 0.0146 |
| Fatty Acid Metabolism | 3 | 2 | 0.0146 |
| Omega-3 fatty acid metabolism | 1 | 1 | 0.0146 |
| Heparan sulfate degradation | 1 | 1 | 0.0146 |
| Fatty acid oxidation, peroxisome | 1 | 1 | 0.0146 |
| Purine metabolism | 1 | 1 | 0.0146 |
| TCA cycle | 1 | 1 | 0.0146 |
| Glycosphingolipid metabolism | 1 | 1 | 0.0146 |

Metabolic Pathways (Mayo Imaging Classification Subclasses)
| Pathway | Metabolites Detected | Detected Metabolites with VIP $\geq 1$ | p-Value<sup>1</sup> |
| --- | --- | --- | --- |
| Porphyrin metabolism | 1 | 1 | 0.0146 |
| Pentose and Glucuronate Interconversions | 1 | 1 | 0.0146 |
| Leukotriene metabolism | 1 | 1 | 0.0146 |
| Phytanic acid peroxisomal oxidation | 1 | 1 | 0.0146 |
| Pentose phosphate pathway | 1 | 1 | 0.0146 |
| Methionine and cysteine metabolism | 3 | 2 | 0.0146 |
| Selenoamino acid metabolism | 1 | 1 | 0.0146 |
| Arginine and Proline Metabolism | 6 | 3 | 0.0295 |
| Xenobiotics metabolism | 4 | 2 | 0.0339 |
| Saturated fatty acids beta-oxidation | 4 | 2 | 0.0339 |
| Glycerophospholipid metabolism | 4 | 2 | 0.0339 |
| Glutamate metabolism | 4 | 2 | 0.0339 |
| Histidine metabolism | 2 | 1 | 0.0458 |
| Tryptophan metabolism | 2 | 1 | 0.0458 |
| Fatty acid activation | 2 | 1 | 0.0458 |
| Drug metabolism - other enzymes | 2 | 1 | 0.0458 |
| Lysine metabolism | 9 | 4 | 0.0469 |
| <sup>1</sup> Gamma-adjusted |  |  |  |

Potentially confounding clinical covariates were further assessed through visualization of OPLS-DA scores plots color-coded by each covariate, which confirmed detected differences in age, hTKV, and mutation distribution across Mayo subclasses (Supplemental Fig. 3).

## Discussion

### Overview

There is currently an unmet need for diagnostic and prognostic indicators of early stages of ADPKD. In this study, we generated LC-MS-based global metabolomic profiles of urine from patients enrolled in the longitudinal EPOC study to identify panels of diagnostic and prognostic metabolic biomarkers of early-stage ADPKD using OPLS-DA models to select metabolite features with the greatest ability to discriminate between control and ADPKD patients (diagnostic) as well as patients with variable rates of disease progression based on MIC subclasses (prognostic). The results of this study showcase the most comprehensive urinary ADPKD metabolomic profile to date and reveal panels of 46 candidate diagnostic biomarkers and 41 candidate prognostic biomarkers. Furthermore, we identified metabolic pathway alterations that may be responsible for early cystogenesis and rapid disease progression that may be potential therapeutic targets.

This work builds on the findings of previous studies that have analyzed urine from ADPKD patients with established disease(9-11). Previous studies have examined metabolic biomarkers in ADPKD patients with a median age of 40.6 years with mean eGFR of 95.5 ml/min per 1.73 m^2^ and hTKV of 1869 ml/m (Gronwald et al.), 46 years with a median eGFR of 62 ml/min per 1.73 m^2^ and hTKV of 935 ml/m (Dekker et al.) and 42 years with median eGFR of 87 ml/min per 1.73 m^2^ and hTKV of 609.9 ml/m^2^ (Hallows et al.)(9-11). In the current study, we selected a substantially younger cohort of individuals with a mean age of 23 years and no evidence of renal impairment, evident by a mean eGFR of 113 ml/min per 1.73 m^2^ and average hTKV of only 491 ml/m. Thus, metabolic markers detected in this younger ADPKD cohort may better reflect early cystogenic pathways that would precede kidney damage and functional decline.

To our knowledge, this is the first study to employ LC-MS to generate urinary metabolomic profiles of ADPKD. Previous studies have employed NMR spectroscopy for metabolomic profiling of urine from ADPKD patients, limiting the analyses to only tens to hundreds of detected metabolites(9, 10). LC-MS offers greater sensitivity than NMR, and thus can detect a wider range of metabolites present at very low concentrations increasing the total number of metabolites detected. Furthermore, LC-MS-based global metabolomic profiling seeks to detect all metabolite features in an unbiased and untargeted fashion, as opposed to focusing on predetermined subsets of known metabolites. Importantly, our approach does not discard data of unidentified metabolite features because identification of these unknown metabolites may result in significant indicators of disease. If mass spectra match to specific known metabolites, putative metabolite identities are reported and mapped to biologically relevant metabolic pathways.

### Comprehensive Global Metabolomic Profile of Early-Stage ADPKD

A comprehensive global metabolomic profile of control and ADPKD cohorts was generated using OPLS-DA, with metabolite features with the greatest ability to discriminate between control and ADPKD cohorts mapped to metabolic pathways. For the OPLS-DA model of control and ADPKD patients, the cumulative 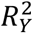 is relatively high and the Q^2^ p-value from the permutation test is small, suggesting that the OPLS-DA model explains a good amount of variation between the cohorts, and there is strong evidence that the model predicts cohort classification better than chance.

The global metabolomic profile of early-stage ADPKD demonstrates profound metabolic reprogramming, with many pathways suggesting a shift from catabolism to anabolism in line with the proliferative potential of ADPKD cells. The global metabolomic profile of early-stage ADPKD was associated with perturbations in pyruvate metabolism, consistent with current literature supporting alterations in energy metabolism in ADPKD. Evidence suggests that ADPKD cells exhibit the “Warburg Effect,” a shift from oxidative phosphorylation to aerobic glycolysis (*i*.*e*., the conversion of pyruvate to lactate)(7). Although aerobic glycolysis generates significantly less ATP per glucose molecule in comparison to oxidative phosphorylation, it supports cell proliferation associated with cyst formation and growth by feeding biosynthetic pathways (*e*.*g*., the PPP) to produce building blocks for organic macromolecules(23).

Furthermore, cell proliferation requires increased synthesis of fatty acids for cell membrane production. Metabolic perturbations we observed in early-stage ADPKD included alterations in a variety of lipid metabolism pathways (fatty acid beta-oxidation, carnitine shuttle, fatty acid metabolism), in line with current evidence suggesting that ADPKD is associated with reduced fatty acid oxidation by inhibiting the carnitine shuttle in favor of increased fatty acid synthesis(7, 24). Proliferation also requires the *de novo* synthesis of amino acids for increased protein synthesis. We identified two amino acid pathways – lysine and tyrosine metabolism – and the urea cycle, which plays a critical role in the *de novo* synthesis of arginine, perturbed in early-stage ADPKD, in support of the changes to protein synthesis required for proliferation for cystogenesis. Furthermore, putative identities of candidate diagnostic and prognostic biomarkers of ADPKD included various urea derivatives. Previous studies that have analyzed the urinary metabolome in ADPKD patients identified metabolite biomarkers in these perturbed metabolic pathways (*e*.*g*., pyruvate, lactate, tyrosine), in further support of our findings(9-11).

A potentially novel finding was the detection of steroid hormone (estrogen and androgen) biosynthesis and metabolism associated with the early-stage ADPKD metabolomic profile. Similarly, putative identities of candidate biomarkers for early detection and prognosis matched to various androgens. Alterations in sex steroid hormone metabolism are linked to sex differences and BMI. However, there were no significant differences in male to female ratios or BMI between control and ADPKD cohorts (Table 1). To further eliminate the possibility that steroid hormone biosynthesis was associated with a potentially confounding clinical co-variate (*e*.*g*., sex, BMI), OPLS-DA scores plots were color-coded to reflect clinical characteristics to showcase that discrimination between cohorts does not reflect sex or BMI (Supplemental Figure 3). The kidneys express both estrogen and androgen receptors, where they play a role in the regulation of calcium and phosphate handling(25). While sex steroid hormone concentrations (androgens and estrogens, in particular) have been previously associated with renal function decline and greater evidence of kidney damage, more data are needed to fully elucidate the underlying mechanism for sex steroid alterations in early-stage ADPKD(26). However, these results suggest that steroid hormones may be candidates for diagnosis and prognostic outcomes for early-stage ADPKD.

### Metabolic Pathway Alterations Associated with Variables Rates of Disease Progression

We also identified metabolic pathways associated with variable rates of disease progression by separating ADPKD patients into MIC subclasses 1A-1E and identifying metabolites with the greatest ability to discriminate between subclasses in OPLS-DA, and mapping metabolites to relevant biological pathways to gain further insight into disease pathogenesis. For the ADPKD OPLS-DA model, the cumulative 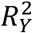 is relatively high, suggesting that the OPLS-DA model explains a good amount of variation between the subclasses. The Q^2^ p-value from the permutation test is not particularly small, so there is inconclusive evidence on the model’s ability to predict subclasses, though this is an expected conclusion due to small sample sizes for each subclass. Furthermore, it is noted that OPLS-DA for more than two classes, as in the case of ADPKD subclasses, introduces some bias when the sample sizes of each class are uneven. Additional validation of this model through analyzing each subclass separately may provide more insight on class separation.

As expected, many metabolic pathways associated with rate of disease progression were similarly identified as a part of the overall metabolomic profile of early-stage ADPKD. This includes pathways involved in energy metabolism (TCA cycle, PPP), steroid hormone biosynthesis and metabolism, fatty acid metabolism (de novo fatty acid biosynthesis, carnitine shuttle, fatty acid oxidation), and amino acid metabolism. These results suggest that these metabolic pathway alterations may be responsible for early cystogenesis and disease heterogeneity.

A number of metabolic pathways that were not detected as a part of the global metabolomic profile of early-stage ADPKD were uniquely associated with the variable rates of disease progression determined by MIC subclasses. These metabolic pathways were involved in ECM remodeling, inflammation, and vitamin D3 metabolism. Metabolic pathways suggestive of ECM remodeling – sialic acid metabolism and chondroitin sulfate and heparan sulfate degradation – were significant in discriminating between ADPKD patients with variable rates of disease progression. Sialic acids, chondroitin sulfates, and heparan sulfates are constituents of proteoglycans which are key components of the ECM basement membranes underlying renal epithelial cells. Early evidence showed that basement membrane remodeling, such as reduced expression of heparan sulfate and chondroitin sulfate, is associated with initial development of cyst formation and is an early feature of ADPKD(27-30). While the ADPKD patients analyzed in the current study have no evidence of renal dysfunction, our results suggest that basement membrane remodeling is an early feature of cyst development that can be detected in the urine of ADPKD patients, and that this may be suggestive of disease prognosis.

Many of the metabolic pathways related to MIC subclasses and disease progression are related to lipid mediators of inflammation, including arachidonic acid, leukotriene, and linoleate metabolism. Putative identities of candidate prognostic biomarkers also included inflammatory lipid mediators eicosapentaenoic acid, linoleic acid, stearolic acid. Inflammation is present in early-stage ADPKD even prior to detectable cysts and subsequent renal function decline, with a linear increase in the levels of inflammatory markers with declining kidney function(31). Arachidonic acid and linoleate metabolism have been previously shown to be significantly perturbed in kidney tissues from a cystic mouse model of PKD(32). These inflammatory lipid mediators are also known to increase cAMP levels to stimulate cell proliferation, fluid secretion, and cyst formation(33, 34). These results suggest that these ADPKD patients may be distinguished from one another based on urinary levels of fatty acid inflammatory markers.

Vitamin D3 metabolism was significantly associated with rate of disease progression determined by the MIC. A recent study investigated the prognostic value of vitamin D3 as a predictor of disease progression and found that 62% of ADPKD patients exhibited vitamin D insufficiency, low levels of serum 25(OH)D and vitamin D receptor (VDR) expression were associated with higher hTKV, and VDR levels negatively correlated with inflammatory markers(35). Studies have also shown that VDR and vitamin D supplementation induce anti-inflammatory actions(36-38). Taken together, these results suggest that alterations in vitamin D3 metabolism may be linked to inflammation early in disease progression and its metabolites may have prognostic value as predictors.

### Panel of Diagnostic and Prognostic Metabolite Biomarkers

From our comprehensive metabolomic profile of early-stage ADPKD, we revealed a panel of 46 metabolite features as candidate diagnostic biomarkers for early detection, and 41 metabolite features associated with disease progression for future validation. It is important to note that candidate metabolite biomarkers are reported according to their mass-to-charge ratios (*m/z* values), because absolute metabolite identification is not obtained by our method. However, putative metabolite identities are reported in-text for metabolite features that matched to metabolite identities with associated KEGG IDs, with the full list of putative identities reported in Supplemental Tables 1 and 3. Unidentified metabolite features were not discarded from the panels because future identification of these unidentified metabolites may result in significant indicators of disease.

The exploratory nature of our study design precludes a causal or mechanistic interpretation of the candidate biomarkers and renal function. However, many of the putative identities of candidate biomarkers for early detection and prognosis are involved in metabolic pathways discussed above (such as urea derivatives, various androgens, inflammatory lipid mediators eicosapentaenoic acid, linoleic acid, stearolic acid) and/or have been previously linked to PKD and/or CKD (such as creatinine, cAMP(39-44), betaine aldehyde(45, 46), phosphoric acid(47-50), and anandamide(51, 52)).

The putative identification of creatinine as a candidate biomarker of early-stage ADPKD was somewhat unexpected because there was no significant difference in plasma creatinine levels between control and ADPKD study participants (Table 1). This result also called into question our selected method for normalization of urinary metabolites. Although it is standard practice to normalize metabolite concentrations to urinary creatinine concentrations to adjust for varying dilutions, the excretion of creatinine is heavily influenced by hydration status, sex, age, diet, muscle mass, and BMI (53, 54). Associated clinical information obtained with control and ADPKD study participants, however, included plasma creatinine concentrations as opposed to urinary creatinine concentrations. We selected PQN as a superior normalization method compared to creatinine adjustment because it was developed specifically for metabolomics research and uses the distribution of metabolites across a pooled QC sample as a normalization factor as opposed to normalizing to a single metabolite(17). Furthermore, a recent study demonstrated PQN was a more reliable method for adjusting urinary concentrations compared to urinary creatinine adjustment(55).

### Limitations

Global metabolomic profiles for control and ADPKD cohorts and ADPKD subclasses were generated using OPLS-DA; however, the OPLS-DA model performance metrics suggest some degree of overfitting and/or poor predictive ability of the models (specifically, for the ADPKD subclasses). This was further limited by the lack of a validation cohort to confirm detected differences in these models. Although control and ADPKD cohorts were very similar based on assessment of clinical characteristics (*e*.*g*., age- and sex-matched), OPLS-DA does not allow for the inclusion of other demographic or clinical covariates, so they must be assessed separately from the model. This study was also limited by relatively small ADPKD patient sample sizes in MIC subclasses when searching for prognostic metabolic biomarkers (1A n=6, 1B n=8, 1C n=15, 1D n=6, 1E n=13). Notably, there were significant differences in clinical characteristics between subclasses (*e*.*g*., age, hTKV, and mutation) which may be potentially confounding covariates and influence detected differences between subclasses. Our global approach revealed metabolite features as candidate biomarkers that were matched to putative identities but did not provide absolute identification. Future studies will further investigate these panels of metabolite features for absolute identification and/or reveal novel metabolites as markers of ADPKD using secondary fragmentation and standards and correlate changes in candidate markers with longitudinal variation in current indicators of renal function (*e*.*g*., eGFR decline).

### Significance and Conclusion

To our knowledge, this pilot study is the first to generate urinary global metabolomic profiles from individuals with early-stage ADPKD with preserved renal function for biomarker discovery. The exploratory dataset reveals metabolic pathway alterations that may be responsible for early cystogenesis and rapid disease progression, and may be potential therapeutic targets and pathway sources for candidate biomarkers. From these results, we generated a panel of candidate diagnostic and prognostic biomarkers of early-stage ADPKD for future validation.

## Acknowledgements

The authors would like to acknowledge the Montana State University Mass Spectrometry CORE Facility for their assistance with mass spectrometry and Statistical Research and Consulting Services for their statistical expertise.

## Grants

Research reported in this publication was supported by the National Institute of General Medical Sciences of the National Institutes of Health under Award Numbers P20GM103474 and R01AR073964 as well as the M.J. Murdock Charitable Trust under Award Number NS-202016444. This study utilized resources provided by the NIDDK sponsored Polycystic Kidney Disease Research Resource Consortium and the Kansas Polycystic Kidney Disease Translation Core Center Award Number U54 DK126126. The content is solely the responsibility of the authors and does not necessarily represent the official views of the funding sources.

## Disclosures

Authors have no conflicts of interest to disclose.

## Author Contributions

EAH, MGG, and AKH designed research experiments; EAH, MGG, ARB, and HDW performed experiments; EAH, MGG, ARB, SKS, MCG, GML, RKJ, and AKH analyzed data; EAH, MGG, ASLY, DPW, RKJ, and AKH interpreted results of experiments; EAH, MGG, SKS, MCG, GML, and AKH prepared figures and tables; EAH, MGG, and AKH drafted manuscript; EAH, MGG, ARB, SKS, HDW, MCG, GML, ASLY, DPW, RKJ, and AKH edited, revised, and approved the final manuscript.

## Notes

### Competing Interest Statement

The authors have declared no competing interest.

