## Supplemental Figures for "Metabolomic Profiling to Identify Early Urinary Biomarkers and Metabolic Pathway Alterations in Autosomal Dominant Polycystic Kidney Disease"

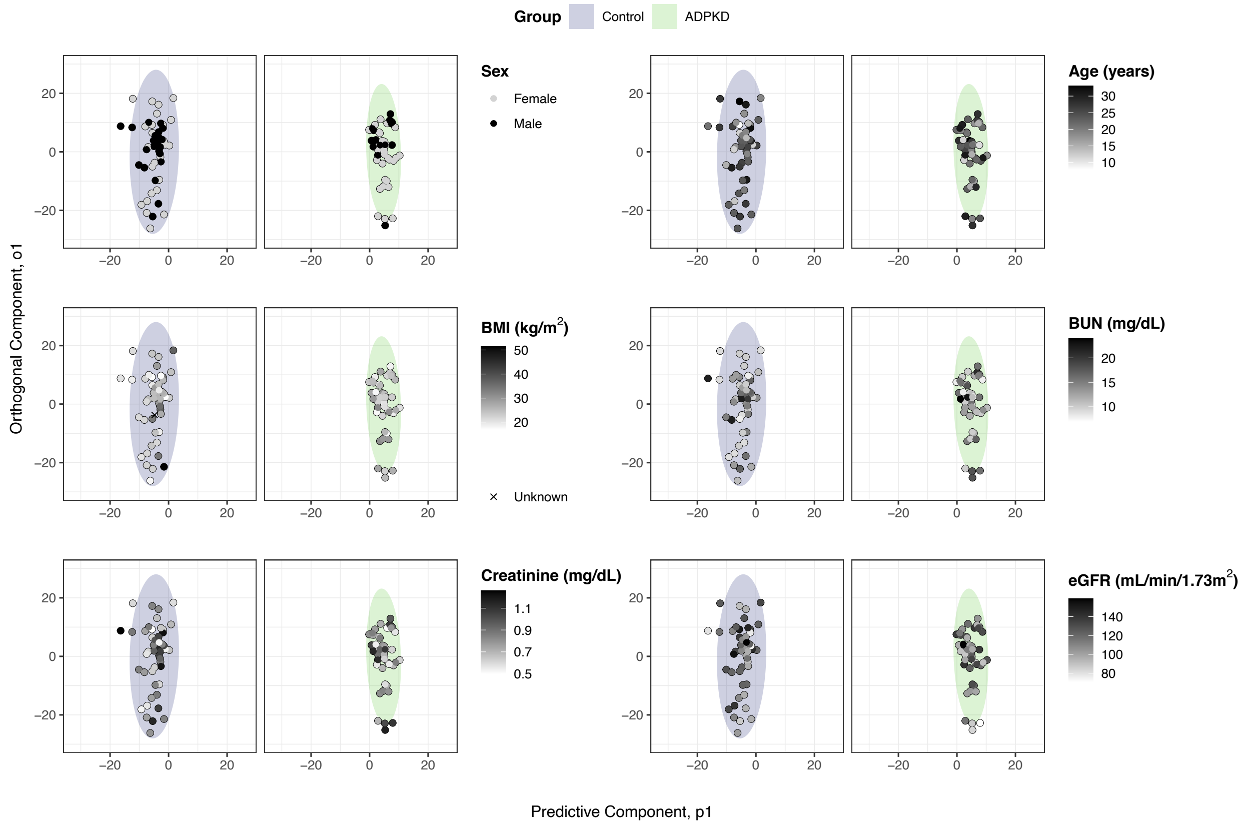


**Supplemental Figure 1. OPLS-DA scores plot of ADPKD and control study participants coded by clinical characteristics.** OPLS-DA scores plot of ADPKD (green) and control (blue) groups from Figure 1 show each subject coded from black to white to reflect potentially confounding clinical covariates. Clinical characteristics visualized included sex, age, BMI, plasma creatinine levels, BUN levels, and eGFR. Cohorts were separated into distinct panels to improve visualization of clinical characteristic differences between cohorts.

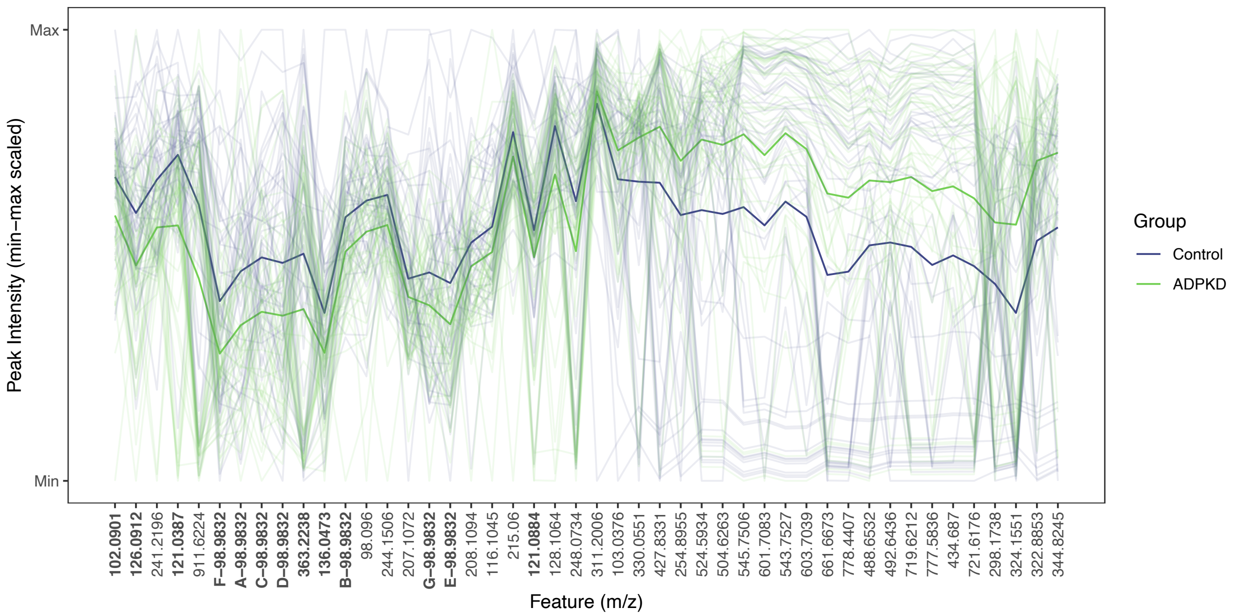


**Supplemental Figure 2. Parallel coordinate plots (PCP) of important metabolite features for discriminating between ADPKD vs. control showcase intragroup variability and subject-level intensity of important features.** PCP of scaled and log transformed peak intensities of important metabolite features selected based on VIP score threshold ≥ 1.9. Features are ordered by hierarchical clustering on group averages. Darker lines indicate mean of each group. Lighter lines indicate individual measurements. Blue represents control cohort peak intensities. Green represents PKD cohort peak intensities. Features named as A-, B-, etc. represent metabolite features with the same *m/z* value but distinct retention times.

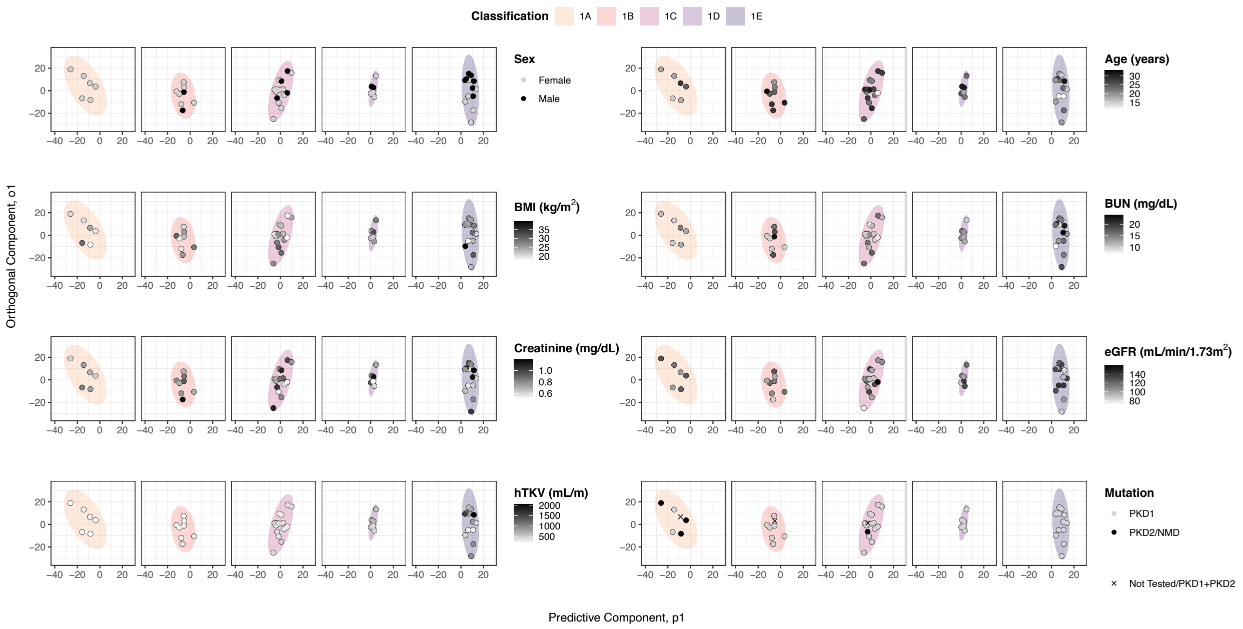


**Supplemental Figure 3. OPLS-DA scores plot of ADPKD study participants classified into MIC subclasses coded by clinical characteristics.** OPLS-DA scores plot of ADPKD MIC subclasses 1A (light orange) to 1E (dark purple) from Figure 3 show each subject coded from black to white to reflect potentially confounding clinical covariates. Clinical characteristics visualized included sex, age, BMI, plasma creatinine levels, BUN levels, eGFR, hTKV, and PKD gene mutation. Cohorts were separated into separate panels to improve visualization of clinical characteristic differences between MIC subclasses. There are noticeable differences and trends in age, hTKV, and mutation between MIC subclasses.

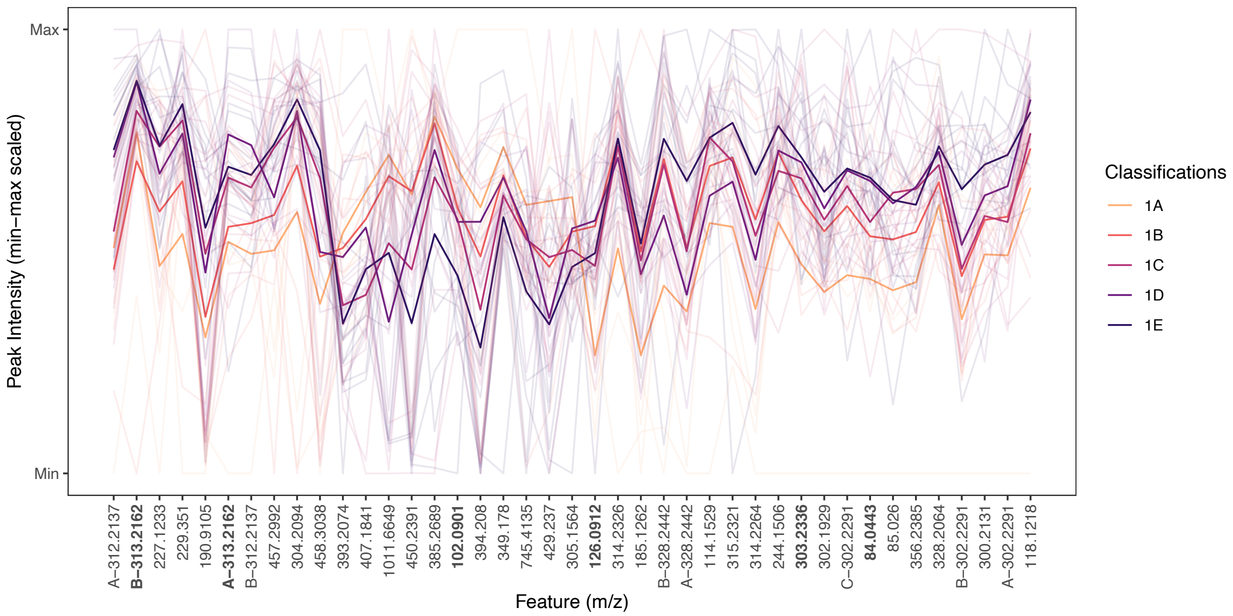


**Supplemental Figure 4. Parallel coordinate plot (PCP) of important metabolite features for discriminating between MIC subclasses showcases intragroup variability and subject-level intensity of important features.** PCP of scaled and log transformed peak intensities of important metabolite features selected based on VIP score threshold ≥ 2. Features are ordered by hierarchical clustering on group averages. Darker lines indicate mean of each group. Lighter lines indicate individual measurements. MIC subclasses are colored by decreasing intensity (1A – light orange; 1E dark purple). Features named as A-, B-, etc. represent metabolite features with the same *m/z* value but distinct retention times.
