## Supplemental Table Captions for "Metabolomic Profiling to Identify Early Urinary Biomarkers and Metabolic Pathway Alterations in Autosomal Dominant Polycystic Kidney Disease"

### Supplemental Tables

**Supplemental Table 1. Candidate diagnostic biomarkers of early ADPKD.** The top discriminatory metabolites selected based on OPLS-DA VIP score threshold ≥ 1.9. Metabolite features are ranked based on VIP score. Features were matched to putative metabolite identities using the metabolite database METLIN (adducts: $H^+$ and $Na^+$, mass tolerance of 30 ppm, excluding drugs, toxins, halogens, peptides). Reported identities are limited to metabolites with KEGG and/or CAS IDs. Metabolite identities are reported with metabolite features *m/z* value and retention time, mass of the putatively identified metabolite, mass accuracy (Δppm), name, chemical formula, CAS and/or KEGG ID, and adduct. Metabolite features may match to multiple putative identities.

**Supplemental Table 2. Full list of pathways altered in early ADPKD based on OPLS-DA VIP scores.** Metabolite features with the greatest ability to discriminate between ADPKD and control study participants in the OPLS-DA (VIP score ≥ 1) were mapped to metabolic pathways using the Functional Analysis module in the metabolomics online program, MetaboAnalyst. Pathways are reported with the total number of metabolites in the pathway (Pathway.total), the total number of metabolite features detected in samples in that pathway (Hit.total), total number of metabolites detected in the pathway with a VIP score ≥ 1 (Hits.sig), Fisher's exact p-value for the pathway (FET), EASE score (modified Fisher's exact p-value) and Gamma-adjusted p-value for the pathway.

**Supplemental Table 3. Candidate prognostic biomarkers to classify ADPKD patients into MIC subclasses.** The top discriminatory metabolites selected based on OPLS-DA VIP score threshold ≥ 2. Metabolite features are ranked according to VIP score. Features were matched to putative metabolite identities using the metabolite database METLIN (adducts: $H^+$ and $Na^+$, mass tolerance of 30 ppm, excluding drugs, toxins, halogens, peptides). Reported identities are limited to metabolites with KEGG and/or CAS IDs. Metabolite identities are reported with metabolite features *m/z* value and retention time, mass of the putatively identified metabolite, mass accuracy (Δppm), name, chemical formula, CAS and/or KEGG ID, and adduct. Metabolite features may match to multiple putative identities.

**Supplemental Table 4. Full list of pathways altered based on rate of disease progression by MIC subclasses.** Metabolite features with the greatest ability to discriminate between MIC subclasses in the OPLS-DA (VIP score ≥ 1) were mapped to metabolic pathways using the Functional Analysis module in the metabolomics online program, MetaboAnalyst. Pathways are reported with the total number of metabolites in the pathway (Pathway.total), the total number of metabolite features detected in samples in that pathway (Hit.total), total number of metabolites detected in the pathway with a VIP score ≥ 1 (Hits.sig), Fisher's exact p-value for the pathway (FET), EASE score (modified Fisher's exact p-value) and Gamma-adjusted p-value for the pathway.
